# Predictive validity of effective shunt fraction in critically ill patients

**DOI:** 10.1101/491167

**Authors:** Emma M Chang, Andrew Bretherick, Gordon B Drummond, J Kenneth Baillie

**Author notes:** These authors contributed equally to this work (Emma M Chang, Andrew Bretherick). Corresponding author: J Kenneth Baillie, The Roslin Institute and Royal (Dick) School of Veterinary Studies, University of Edinburgh, Easter Bush, EH25 9RG, UK., 0131 651 9100.

## Abstract

Accurate measurement of pulmonary oxygenation is important for classification of disease severity and quantification of outcomes in clinical studies. We compared predictive validity of established tension-based methods with two new measures of shunt fraction: (1) a non-invasive effective shunt (ES); and (2) inferred values from an integrated mathematical model of gas exchange (DB). Median absolute error (MAE) values for the four measures considered were: alveolar-arterial difference, 7.30kPa; P_a_O_2_/F_I_O_2_ ratio, 2.41kPa; DB, 2.13kPa; ES: 1.88kPa. ES performed significantly better than other measures (p<10^−10^ in all comparisons). While the simplicity of P/F is suitable for routine use, the superior predictive validity of ES should make this measure the preferred choice where physiological accuracy is important, such as for use as surrogate outcome in clinical research.

## Introduction

Hypoxia is the defining feature of respiratory failure. Accurate quantification of pulmonary oxygenation defect is essential to determine inclusion in clinical trials, measure outcomes in research studies, and to observe change in lung function in a clinical setting.

In severely hypoxic patients, direct measurement of intrapulmonary shunt provides the most accurate quantification of an oxygenation defect [1]. Tension-based indices, including P_a_O_2_/F_I_O_2_ (P/F) ratio and alveolar - arterial (A-a) difference have poor agreement with intrapulmonary shunt fraction [1–3]. The primary limitation in tension-based indices is the marked and non-linear change in P_a_O_2_ when F_I_O_2_ is changed [4]. Brochard and colleagues demonstrated that this can be predicted from a simple mathematical model[5].

The concept of predictive validity is a mathematical reality check for a clinical measure. For a given clinical measure, predictive validity quantifies the extent to which that measure predicts an unseen event. This enables a rigorous, unbiased test of how well a clinical measure is describing a real entity. This approach, using mortality as the predicted event, was used in the development of consensus definitions for both acute respiratory distress syndrome (ARDS)[6] and sepsis [7].

A measure that accurately reflects the true state of a patient’s lungs should not change markedly following a change in F_I_O_2_. Therefore, the prediction of a P_a_O_2_ following a change in F_I_O_2_, assuming that the measure of the oxygenation defect remained unaltered, is a valid assessment for predictive validity.

We hypothesised that an easily-understood, content-based oxygenation index may be obtainable from routinely-acquired arterial blood gas (ABG) data, without any need for additional invasive measurements. In order to assess different approaches, we quantified the predictive validity of P/F, A-a and two new methods of estimating shunt fraction (effective shunt fraction (ES) and a database method (DB)) in a simple test: prediction of P_a_O_2_ following a change in F_I_O_2_ in a large retrospective cohort.

## Materials and Methods

### Data source and filtering

We used a set of 78,159 arterial blood gas samples taken from patients on the intensive care unit (ICU) at the Royal Infirmary of Edinburgh between 2011 and 2016. This contained data from 6,511 patients. To obtain pairs of ABGs in which underlying pulmonary pathology was unlikely to change substantially between samples, we limited the analysis to pairs of samples taken within a 3hr window from weaning patients (F_I_O_2_ decreasing) with stable alveolar ventilation (change in P_a_CO_2_ < 0.3kPa). A baseline measure, reflecting the expected noise in predictive validity estimates, was calculated from pairs of ABGs meeting these criteria, but in which F_I_O_2_ remained constant.

### Predictive validity

Each measure was quantified for the earlier ABG in each pair. For ES, P/F and A-a, the F_I_O_2_ in the second ABG was used to estimate the paired P_a_O_2_. For the DB method, input settings for an integrated mathematical model of gas exchange were identified which matched to the first ABG. These were then extrapolated to the F_I_O_2_ and P_a_CO_2_ of the second ABG, and the mean P_a_O_2_ of all matching model runs taken to be the prediction (Figure 1b). Median absolute differences between predicted and observed P_a_O_2_ across all ABG pairs were taken as the predictive validity for each method.

**Figure 1:**
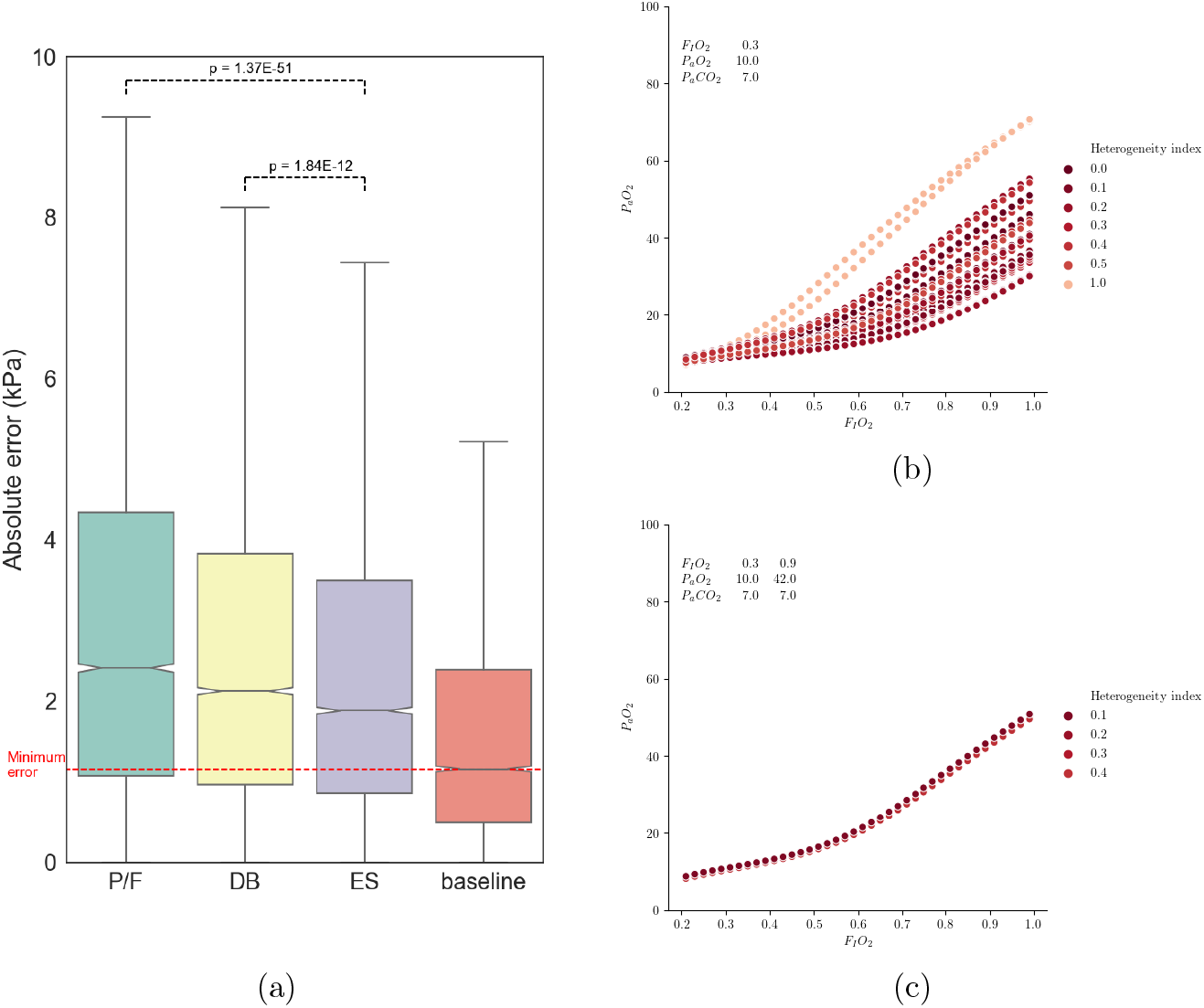
(a) Boxplot showing distribution of absolute error for each measure in all samples, together with baseline distribution of pairs of ABGs in which F_I_O_2_ was unchanged. (b) Range of possible F_I_O_2_-P_a_O_2_ combinations for conditions matching a single ABG result. (c) Range of possible results for conditions matching two ABG result at different F_I_O_2_.

### Research Ethics

Ethical approval was obtained from the Scotland A Research Ethics Committee [16/SS/0209].

### Software and statistical analyses

All analysis was performed using Python 3.5.2 and scipy.stats version 0.18.1. A Kruskal-Wallis H-test was used to determine the difference in error rate between the different measures. Wilcoxon signed-rank tests with Bonferroni correction were used as a stringent post test for pairwise comparisons.

## Results

### Comparison of oxygenation measures

From a total of 78159 ABGs from 6511 patients, a test set was selected at random, containing 54115 ABGs from 4558 patients. In a formal test set comprising 9635 pairs of ABGs, median absolute error (MAE) values for the four measures considered were: A-a, 7.30kPa; P/F, 2.41kPa; DB, 2.13kPa; ES: 1.88kPa. ES had significantly superior predictive validity than all other measures Table 1.

**Table 1:** Pairwise comparisons between errors in oxygenation measures in test set (Mann-Whitney u test, Bonferroni correction).

| Measure 1 | Measure 2 | p (MWu) | p Threshold | Reject null |
| --- | --- | --- | --- | --- |
| A-a | P/F | <1E-300 | 0.008 | True |
| A-a | DB | <1E-300 | 0.008 | True |
| A-a | ES | <1E-300 | 0.008 | True |
| P/F | DB | 6.86E-17 | 0.008 | True |
| P/F | ES | 1.37E-51 | 0.008 | True |
| DB | ES | 1.84E-12 | 0.008 | True |

### Validation of assumed values

Three key assumed values are required for the calculation of ES: respiratory exchange ratio (RER), cardiac output (Q) and metabolic oxygen consumption (VO_2_). Varying these values over wide ranges (RER, 0.8 to 1.1; Q, 3 to 15 l.min^-1^; VO_2_, 0.15 to 1 l.min^-1^) had minimal effect on the value of effective shunt.

## Discussion

This is the first study, to our knowledge, comparing the predictive validity of non-invasive effective shunt fraction with tension-based measures. Our observations are consistent in magnitude and direction with previous work using direct measurements of pulmonary shunt in human participants [1–3].

The poor performance of A-a difference is consistent with the report by Cane and colleagues [1], who demonstrated that A-a was the least reliable measure compared with invasive measurements of Qs/Qt. The very poor predictive validity of A-a in our study, together with previous work, leads us to conclude that this measure has no role in any context.

Marked changes in PaO2 may occur within a 3h interval due to real changes in pulmonary function, for example due to recruitment, suction, diuresis, or change in posture. We therefore cannot draw any inference from the absolute value of the error in prediction, only a comparison between different methods. Noise caused by these and other factors is expected to limit the maximum possible accuracy of any prediction of P_a_O_2_. This minumum achievable error is reflected by the baseline values (Figure 1a) showing the change in P_a_O_2_ between pairs of ABGs meeting the other selection criteria, with no change in F_I_O_2_.

Since V/Q heterogeneity and shunt are separate inputs into the physiological model used to generate the database (see Supplementary Information), this approach is expected to handle this distinction better than the other measures. However, as shown in Figure 1b, there is insufficient information in a single ABG to distinguish between shunt and V/Q heterogeneity. In contrast, with two ABGs taken at different settings of F_I_O_2_, the patient’s oxygen responsiveness is quantified, greatly restricting the range of possible values for both shunt and V/Q heterogeneity (Figure 1c). This double F_I_O_2_ test may resolve the uncertainty in quantifying pulmonary shunt, but is computationally-intensive.

The effective shunt fraction can be calculated on any ABG result, provided the F_I_O_2_ is known. The computation is fast and simple. Hence the method could be retrospectively applied to previous studies that hold ABG data in a machine-readable format. Whilst the simplicity of the P/F ratio will continue to make it the best choice for clinical use, the superior predictive validity of ES makes it a better choice where accurate quantification of oxygenation defect is necessary.

## Supporting information

## Code availability

An online calculator to compute the effective shunt fraction is available at: http://baillielab.net/es

Python code to calculate the effective shunt fraction is available from github: http://github.com/baillielab

## Author contributions

E.M.C. and J.K.B. designed the study and conducted computational analysis. J.K.B. and A.B wrote the computational model of gas exchange. G.B.D. contributed to the conception of hypothesis and interpretation of the results. J.K.B., E.M.C. and A.B. wrote the manuscript with assistance from G.B.D. All authors commented on or contributed to the final manuscript.

