## Supplementary material for "Predictive validity of effective shunt fraction in critically ill patients"

### Design of an integrated model of gas exchange

#### Methods

We designed a simple, integrated physiological model based on standard physiological relationships Figure 1. We followed the principle that the mass balance of the lung, for oxygen and carbon dioxide, must be preserved - the amount of oxygen leaving (or carbon dioxide entering) the alveolar space per minute must equal the amount added (or removed) from the blood during that time[1].

At steady state, the amount of oxygen removed from the alveolar space, and added to the blood, must also equal that consumed by metabolism; the converse is true for carbon dioxide. These values are dictated by metabolism and are linked by the respiratory quotient (Equation 1).

Equation 2 symbolically describes the mass balance of oxygen in the alveolar space; Equation 3 the mass balance of carbon dioxide. Equations 4 and 5 are the equivalent equations in the blood phase.

$$\dot{V}_{C_{O2}} = \dot{V}_{O_2}.RQ \tag{1}$$

$$\dot{V}_{O_2} = F_{I_{O_2}}.\dot{V}_I - F_{A_{O_2}}(\dot{V}_I - \dot{V}_{O_2} + \dot{V}_{CO_2}) \tag{2}$$

$$\dot{V}_{CO_2} = F_{A_{CO_2}}(\dot{V}_I - \dot{V}_{O_2} + \dot{V}_{CO_2}) - F_{I_{CO_2}}.\dot{V}_I \tag{3}$$

$$\dot{V}_{O_2} = (C_{c'_{O_2}} - C_{\bar{v}_{O_2}})(\dot{Q}_t - \dot{Q}_s) \tag{4}$$

$$\dot{V}_{CO_2} = (C_{\bar{v}_{CO_2}} - C_{c'_{CO_2}})(\dot{Q}_t - \dot{Q}_s) \tag{5}$$

The model performs in the blood (using Equation 4 and Equation 5) an analogous calculation to that performed in the alveolar space (using Equation 2 and Equation 3) by the alveolar gas equation.

For simplicity and computational speed, we used a three compartment (dead-space, shunt fraction and a single exchanging lung compartment) steady-state

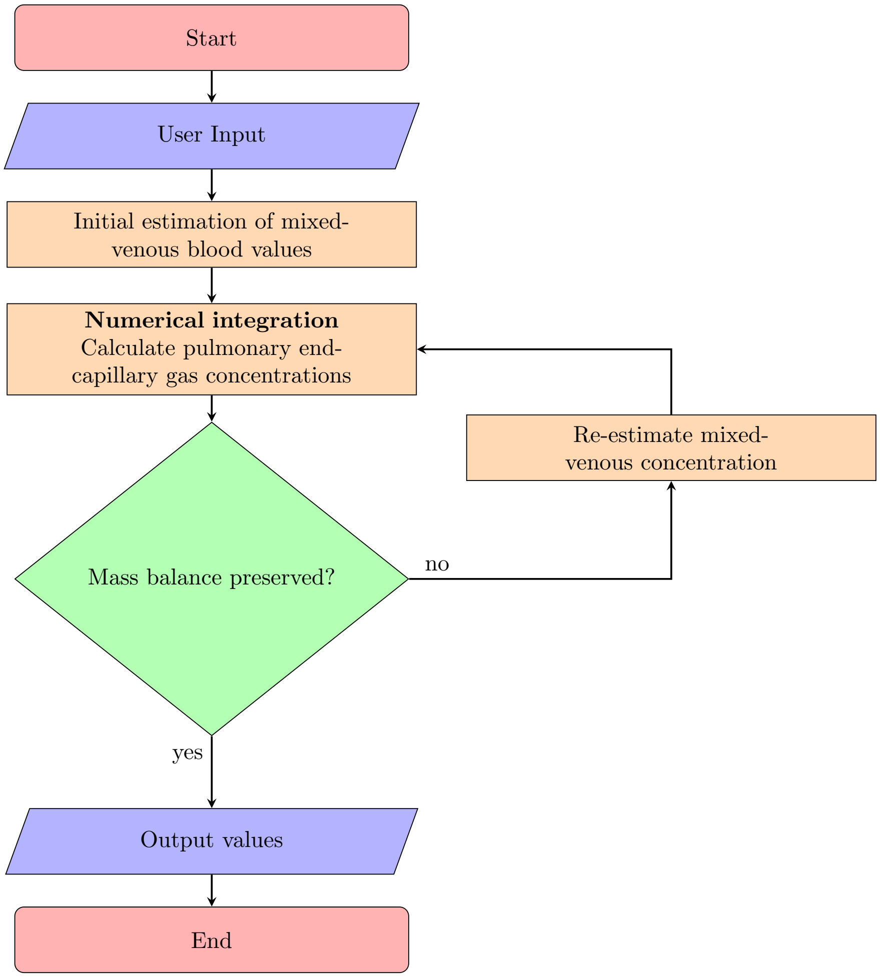


Figure 1: Structure of model. The model reads the user input and takes an initial estimate of mixed-venous values based on the alveolar gas equation. It then performs a modified Bohr integral using a method similar to that of Wagner & West[2] to determine the associated pulmonary end-capillary values. The amount of oxygen leaving (or carbon dioxide entering) the alveolar space per minute must equal the amount added (or removed) from the blood during that time. The model iterates to find a solution that achieves this before displaying final output values.

model. Dead space is the volume of each tidal volume breath that does not reach parts of the lung participating in gas exchange; shunt fraction as the analogous, but hypothetical, fraction of blood that passes through the lungs without participating in gas exchange. Alveolar ventilation is determined by respiratory rate, tidal volume and dead-space, lung perfusion by cardiac output and shunt fraction - all determined by the user.

##### Inter-conversion between whole blood content and partial pressures

We employed the formulae of Dash and Bassingthwaighte[3], with minor correction and modifications, and their analytical or numeric inversions as appropriate.

$$C_{n_{O_2}} = ([O_2] + 4.[Hb].S_{n_{O_2}})\frac{R.T}{P} \tag{6}$$

$$C_{n_{CO_2}} = ([CO_2] + 4.[Hb].S_{n_{CO_2}} + [HCO_3^-])\frac{R.T}{P} \tag{7}$$

Whole blood content of oxygen and carbon dioxide were expressed as functions of their respective partial pressures, erythrocyte pH and the $P_{50}$ (oxygen) of haemoglobin. Unfortunately, the equations for whole blood content are not amenable to analytical inversion. We obtain numeric solutions using root-finding algorithms.

The Henderson-Hasselbalch equation[???] describes the relationship between pH, acid and salt and can be applied to the bicarbonate buffer system that operates in mammalian blood (Equation 8).

$$pH = pK_a + log_{10}\left(\frac{[HCO_3^-]}{\alpha_{O_2}.P_{a_{O_2}}}\right) \tag{8}$$

The van Slyke equation describes the carbon dioxide equilibration curve of blood *in vitro*. Simultaneous solution of the van Slyke and the Henderson-Hasselbalch equations allows for estimation of acid-base state where only one of bicarbonate concentration, partial pressure of carbon dioxide or pH is known. We made use of user-determined base excess, in fully saturated blood, and applied the van Slyke equation as described by Lang & Zander [Lang W, Zander R: The accuracy of calculated base excess in blood. Clin Chem Lab Med 2002, 40:404-410.].

##### Diffusion limitation

We modelled diffusion limitation of oxygen in the gas exchanging lung compartment by taking the initial conditions of blood entering the lungs as those of mixed-venous blood. Using a method similar in nature to that of Wagner &

West[2], a set of differential equations describing alveolar gas and pulmonary capillary blood were obtained and solved numerically. In contrast with the method of Wagner and West, we allowed alveolar gas composition to vary with time (as gases are exchanged with the blood) and made the simplifying assumption that the rate of reaction of carbon dioxide (as a whole) with blood is instantaneous. The complex interaction of carbon dioxide and blood was not modelled and for this reason diffusion limitation of carbon dioxide transfer is not discussed further.

Equation 9 and Equation 10 represent metabolic consumption of oxygen and production of carbon dioxide as functions of mixed-venous oxygen and carbon dioxide concentration. We solve this set of simultaneous equations (Figure 2), using numerous inter-conversions between whole blood content and partial pressure, each time an input value changes.

$$\begin{aligned} V_{O_2} &= (C_{c'_{O_2}} - C_{\bar{v}_{O_2}}) \times (\dot{Q}_t - \dot{Q}_s) \\ V_{O_2} &= (f(C_{\bar{v}_{O_2}}, C_{\bar{v}_{CO_2}}) - C_{\bar{v}_{O_2}}) \times (\dot{Q}_t - \dot{Q}_s) \end{aligned} \tag{9}$$

$$\begin{aligned} V_{CO_2} &= (C_{\bar{v}_{O_2}} - C_{c'_{CO_2}}) \times (\dot{Q}_t - \dot{Q}_s) \\ V_{CO_2} &= (C_{\bar{v}_{O_2}} - g(C_{\bar{v}_{O_2}}, C_{\bar{v}_{CO_2}}) - C_{\bar{v}_{O_2}}) \times (\dot{Q}_t - \dot{Q}_s) \end{aligned} \tag{10}$$

#### Abbreviations

Symbols are constructed in three parts: a primary symbol and a two part subscript.

The primary symbol describers what quantity the symbol is in relation to; the secondary part, where it is in relation to and the tertiary part, what substance.

##### Primary symbol

C concentration of gas in blood L / L

F fractional concentration

P pressure *or* partial pressure kPa

blood flow (volume per time) L / min

RQ respiratory quotient fraction

S haemoglobin saturation fraction

T temperature Kelvin

gas flow (volume per time) L / min

$\alpha$

solubility co-efficient M.kPa$^{-1}$

**Secondary symbols**

**Gas phase**

A alveolar

I inspired

**Blood phase**

pulmonary end-capillary

mixed-venous

n *n*th vascular compartment

**Other**

s pulmonary shunt

t total blood flow; cardiac output

**Substance**

$CO_2$ carbon dioxide

Hb haemoglobin (tetrameric)

$HCO_3^-$ bicarbonate

$O_2$ oxygen

**Other symbols**

[ $x$ ] concentration of specified substance, $x$ M

$P_{50(O2)}$ partial pressure of oxygen at which

oxygen-haemoglobin saturation is 50% kPa

R universal gas constant

#### Tables

Table 1: Model inputs and suggested 'standard' values.

| **Respiratory** | **Cardiovascular** | **Haematological** | **Metabolic** |
|---|---|---|---|
| Respiratory rate *12 breaths / min* | Cardiac output *6.5 L / min* | Haemoglobin concentration *150 g / L* | $V_{O2}$ *0.250 L / min* |
| Tidal volume *0.475 L* | Pulmonary shunt fraction *0.02* | Base excess *0 mEq / L* | Body temperature *310.15K; 37 °C* |
| Dead space volume *0.110 L* | $V_c$ *0.075 L* | 2,3-DPG concentration *0.00465 M* | Tissue shunt fraction *0.05* |
| $F_{IO2}$ *0.21* | | Haematocrit *0.45* | Respiratory quotient *0.8* |
| Altitude *0 m* | | | |
| $D_{mO2}$ *0.3 L / (min . kPa)* | | | |

These variables represent the modifiable inputs that the user is able to adjust in this on-line model and their default values when the user initially enters the web site.

$F_{IO2}$, fraction of inspired oxygen; $D_{mO2}$, diffusing capacity of the alveolar-membrane for oxygen; $V_c$, pulmonary capillary volume; 2,3-DPG, 2,3-diphosphoglycerate; $V_{O2}$, minute consumption of oxygen.
