## Supplementary material for "Predictive validity of effective shunt fraction in critically ill patients"

### Supplementary information

#### Derivation of oxygenation measures and prediction of $P_aO_2$

##### Estimated Shunt Fraction

The estimated shunt fraction was calculated by assuming the necessary values to complete the shunt equation:

$$\frac{Q_S}{Q_T} = \frac{C_cO_2 - C_aO_2}{C_cO_2 - C_vO_2} \tag{1}$$

An estimation of the shunt fraction (Qs/Qt) was calculated using a combination of Equation 1 and the direct Fick equation to form Equation 2. Arterial (C_a$O_2$) and end-capillary ($C_{c'}O_2$) oxygen contents were derived using model equations from Dash and Bassingthwaighte to estimate the $PO_2$ at which Hb is 50% saturated ($P_{50}$). Combining this value with $P_aO_2$ allows $C_aO_2$ to be obtained. $C_{c'}O_2$ was estimated using $P_AO_2$ from the alveolar gas equation (Equation 9). The combination of the two equations on the left provides Equation 10 from which we obtained $C_vO_2$.

Values for oxygen consumption ($VO_2$) and cardiac output (Q) were set at single optimal values in the physiolgical range (see below).

$$C_vO_2 = C_aO_2 \times \frac{DO_2 - VO_2}{DO_2}$$

$$DO2 = C_aO_2 * Q$$

$$C_vO_2 = C_aO_2 - \frac{VO_2}{Q}$$

$$\frac{Q_S}{Q_T} = \frac{C_cO_2 - C_aO_2}{C_cO_2 - C_aO_2 - \frac{VO_2}{Q}} \tag{2}$$

By utilizing an expanded version of Equation 1 the following expression was formed to estimate the 'new' C_a$O_2$ in ABG2:

$$C_aO_2 = C_cO_2 - \frac{Q_S/Q_T \times VO_2}{Q_T - Q_S} \tag{3}$$

Derivation of the Estimated Shunt equation from the expanded shunt fraction:

$$Q_S(C_CO_2 - C_vO_2) = Q_T(C_cO_2 - C_aO_2)$$

$$Q_TC_aO_2 = Q_TC_cO_2 - Q_SC_cO_2 + Q_SC_vO_2$$

$$Q_TC_aO_2 = Q_TC_cO_2 - Q_SC_cO_2 + Q_S\left(\frac{Q_TC_aO_2 - VO_2}{Q_T}\right)$$

$$Q_TC_aO_2 = Q_TC_cO_2 - Q_SC_cO_2 + Q_SC_aO_2 - \frac{Q_SVO_2}{Q_T}$$

$$(Q_T - Q_S)C_aO_2 = (Q_T - Q_S)C_cO_2 - (Q_S/Q_T \times VO_2)$$

$$C_aO_2 = C_cO_2 - \frac{Q_S/Q_T \times VO_2}{Q_T - Q_S} \tag{4}$$

The corrected version of Dash and Bassingthwaighte's work[1] provides simple mathematical expressions based on the nonlinear biochemical interactions of $O_2$ and $CO_2$ with Hb. The invertible equation contains 'binding constants' ($KHb_{O2}$) that depend on $P_aO_2$ among other factors. By reversing the equation, our $P_{50}$ function can predict $P_aO_2$ from Hb saturation and vice-versa.

An online calculator to compute the effective shunt fraction is available at:

http://baillielab.net/es

Python code to calculate the effective shunt fraction is available from github:

http://github.com/baillielab

#### P/F Ratio

P/F was first calculated using the $P_aO_2$ and corresponding $F_IO_2$ from ABG1. By rearranging Equation 5 the new $P_aO_2$ can be predicted using the original P/F value with the new $F_IO_2$ value from ABG2 via Equation 6.

$$P/F = \frac{P_aO_2}{F_IO_2} \tag{5}$$

$$P_aO_2 = P/F \times F_IO_2 \tag{6}$$

#### Alveolar-arterial difference:

The equation for Alveolar-arterial difference is shown below:

$$Aa\,difference = P_AO_2 - P_aO_2 \tag{7}$$

Therefore Equation 8 was used to estimate $P_aO_2$ following a change in $F_IO_2$.

$$P_aO_2 = P_AO_2 - Aa\,difference \tag{8}$$

Whereby:

$$P_AO_2 = P_IO_2 - \frac{P_aCO_2}{RER} + \left[F_IO_2 \times P_aCO_2 \times \left(\frac{1 - RER}{RER}\right)\right] \tag{9}$$

Where:

$$DO_2 = Q \times C_aO_2$$

and

$$C_vO_2 = \frac{(DO_2 - VO_2)}{Q}$$

$$VO_2 = Q \times (C_aO_2 - C_vO_2) \tag{10}$$

Alveolar oxygen partial pressure ($P_AO_2$) was calculated using values from ABG1 to obtain the A-a difference, then calculated again with values from ABG2 to predict $P_aO_2$ in Equation 8. RER was kept constant at 0.8 for ES and A-a difference.

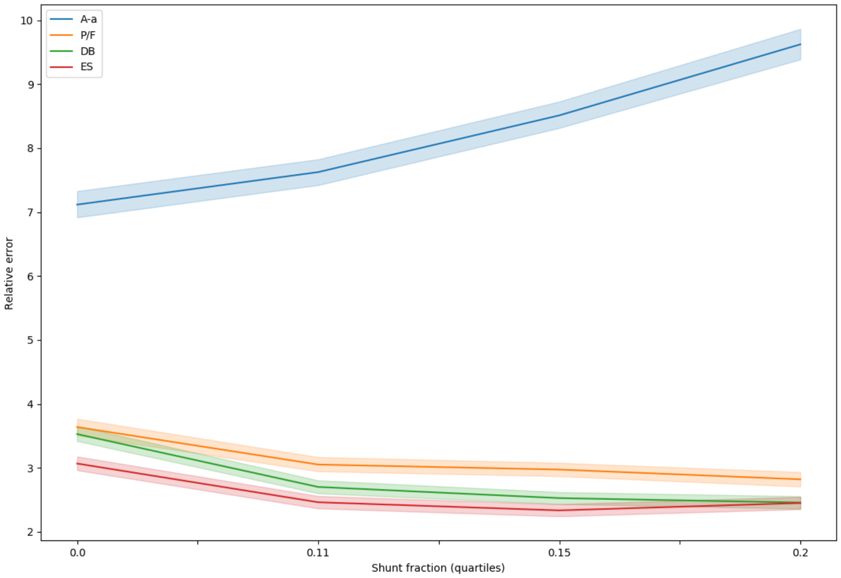


Figure 1: Change in relative error (absolute error / $P_aO$~2 in second ABG) for each measure across shunt severity quartiles.

### Optimisation of assumed variables

The test set of ABGs was used for optimisation of the variables listed below. Although VO2 and Q have a direct impact on the value of ES (Equation 2), when these values are held constant they have no impact on the accuracy of predictions made using ES (Figure 3 and Figure 4).

#### Respiratory exchange ratio (RER)

Both the calculation of the $C_{c'}O_2$ term in Equation 2 for effective shunt fraction (ES) and the alveolar term in A-a difference require the alveolar gas equation, Equation 9, which uses RER to calculate $P_AO_2$. We therefore considered the possibility that changing the assumed RER would affect the predictive validity of these measures.

Varying the assumed RER across the range 0.7 to 1.2 had negligible effect on ES and a slight effect on A-a (Figure 2). We therefore chose a value of 1 for

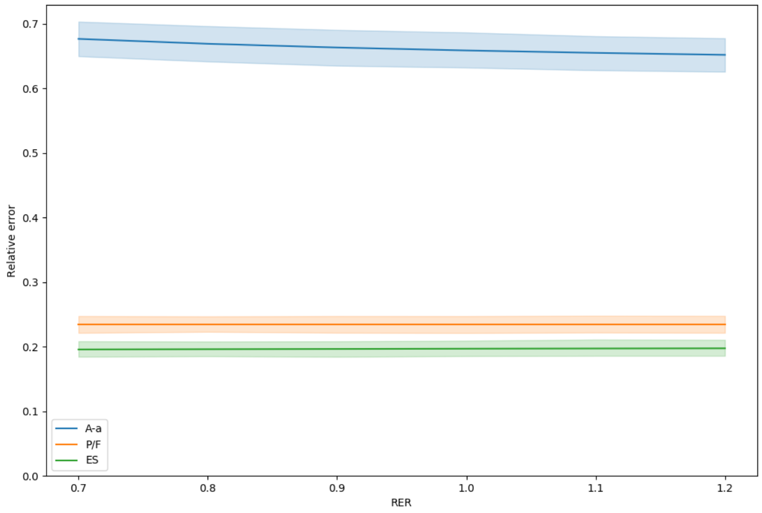


Figure 2: Change in relative error (absolute error / $P_aO$~2 in second ABG) of P/F, ES, and A-a across a range of assumed values for RER

#### Cardiac output (Q)

Changing cardiac output across the range 1 to 15 $l.min^{-1}$ had minimal effect on the accuracy of predictions by ES (Figure 3)

#### Metabolic oxygen consumption (VO2)

Likewise, a change in VO2 does not alter the predictive validity of ES.

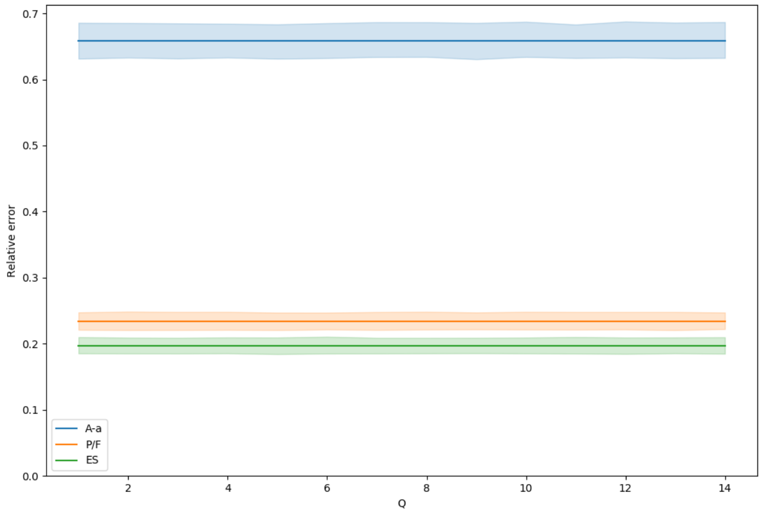


Figure 3: Change in relative error (absolute error / $P_aO$~2 in second ABG) of P/F, ES, and A-a across a range of assumed values for Q

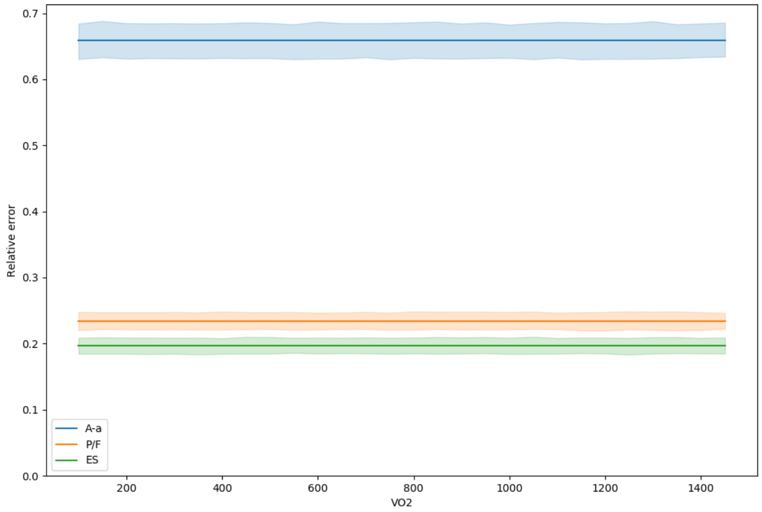


Figure 4: Change in relative error (absolute error / $P_aO$~2 in second ABG) of P/F, ES, and A-a across a range of assumed values for Q
